# Is shorter faster? The geometry and aerodynamics of crank length in cycling

**DOI:** 10.64898/2026.09.25.754230

**Authors:** Scott L. Delp, Ellen Kuhl

## Abstract

Crank length affects the geometry of the bicycle–rider system and influences the motion of the lower limb throughout the pedal cycle. Conventional crank selection has largely relied on rider stature and mechanical power, yet mechanical power and cycling economy remain remarkably insensitive to crank length across a broad range. Yet, elite cyclists and triathletes increasingly adopt shorter cranks, particularly in disciplines where aerodynamic drag strongly affects cycling performance. However, the mechanisms that connect crank length to mechanical power and aerodynamic drag remain unclear. Here we show that crank length couples rider anthropometry, hip kinematics, and aerodynamic drag. Population anthropometry of 6,068 individuals reveals that stature alone poorly predicts personalized crank length, while geometric analysis identifies a nearly linear relationship between crank length and minimum lower hip angle: a 10-mm reduction in crank length increases hip-angle margin by approximately 1.42° with little mechanical penalty. This geometric freedom permits more flexion of the torso and a more aerodynamic posture; the resulting aerodynamic advantage grows rapidly with cycling speed because aerodynamic power scales with velocity cubed. Data from 150 elite cyclists and triathletes support this shift toward shorter cranks, with triathletes adopting shorter relative crank lengths than cyclists. Our results establish crank length as a personalized design variable that connects lower-limb geometry to aerodynamics. These insights enable personalized crank selection, integrated rider–bicycle design, and performance optimization for the increased aerodynamic demands of high-speed cycling.

## 1. Introduction

### Crank length presents a long-standing paradox in cycling biomechanics

It changes pedaling geometry substantially, yet it changes mechanical power and cycling economy surprisingly little [12]. Maximal cycling power remains remarkably insensitive to crank length across a broad range, with substantial losses only for unusually short or long cranks [15]. Similarly, small changes in conventional crank lengths have little effect on gross efficiency, heart rate, or metabolic cost during submaximal cycling [5]. Recent experiments with 165-, 170-, and 175-mm cranks found no significant differences in cycling economy or sprint power in trained cyclists [13]. Together, these studies identify a mechanically and metabolically neutral regime in which cyclists can alter crank length with little mechanical or metabolic consequence.

### Crank length strongly alters rider geometry

Longer cranks increase hip and knee angular excursions and alter joint angular velocities, whereas ankle kinematics remain comparatively insensitive to crank length [2]. Crank length can also alter joint-specific mechanics under fixed-cadence conditions even when total external power changes little [1]. Recent three-dimensional motion analysis confirms that small reductions in crank length reduce hip and knee flexion without a detectable change in mean power [16]. These effects depend not only on stature, but also on the individual proportions of the thigh, shank, and foot. A crank length that produces one configuration in one rider can produce a different configuration in another rider of similar stature. This distinction challenges simple crank-to-stature scaling and places *individual geometry* at the center of crank-length selection.

### Geometry links crank length to aerodynamics

This geometric effect may matter more for cycling performance than the direct effect of crank length on mechanical power. Aerodynamic drag dominates resistive power demand at racing speeds [17], and small changes in torso position can substantially alter aerodynamic drag area [6]. A shorter crank reduces the pedal-circle radius and decreases hip flexion near the top of the pedal stroke. The resulting *hip-angle margin* can permit a lower torso position without further hip closure. Previous studies have therefore established two separate relationships: crank length alters lower-limb kinematics [2], and torso position alters aerodynamic drag [17]. The *quantitative connection* between these relationships, however, remains unresolved. This gap becomes particularly important in long-course triathlon, where athletes sustain high speeds for several hours without drafting.

Here we treat crank length not primarily as a power-production parameter, but as a *geometric design variable* that couples rider anthropometry, joint kinematics, bicycle position, and aerodynamic drag. We first establish lower-limb scaling across 6,068 individuals and quantify the variation that stature alone cannot explain. We then determine how crank length and individual segment geometry control hip and knee kinematics throughout the pedal cycle and quantify the sensitivity of minimum lower hip angle to crank length. We combine this geometric scaling with experimental crank-length–power relationships and aerodynamic measurements to connect crank length to drag area and net power change. Finally, we compare these predictions with crank-length choices in 150 elite cyclists and triathletes. *We hypothesize that crank length acts primarily through geometry rather than mechanical power, and that the influence of crank length on net power balance increases with cycling speed*.

## 2. Methods

### 2.1. Population anthropometry

We quantify lower-limb geometry using population-scale anthropometric data from the ANSUR II database, with 4,082 men and 1,986 women [8]. We define thigh length *L*_thigh_ as the difference between trochanterion height and lateral femoral epicondyle height and shank length *L*_shank_ as the difference between lateral femoral epicondyle height and lateral malleolus height. We define leg length as

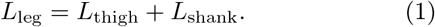

We quantify sex-specific population distributions and fit proportional relationships of the form *L*_*i*_ = *m*_*i*_*H* through the origin, where *H* denotes stature and *i* ∈ *{* thigh, shank, foot *}*. We use all available individuals for each fit and restrict the displayed scaling relationships to the 1st–99th percentile range of stature. To quantify variation among individuals of similar stature, we also compare thigh and shank lengths within *±*1 cm of the sex-specific mean stature.

### 2.2. Cycling kinematics

We construct a planar geometric model of the rider–bicycle system (Fig. 1). The reference configuration uses a crank length *L*_c_ = 170 mm, a saddle height *L*_h_ = 70 cm, and a saddle setback *L*_s_ = 5 cm behind the bottom bracket. We position the hip joint 6 cm behind and 2 cm above the saddle tip, which is located at a distance *d* from the center of the bottom bracket. We represent the pedal-to-ankle segment with a reference length of 17 cm and scale this length in direct proportion to anthropometric foot length. We vary the foot angle smoothly from 28° at top to 43° at bottom dead center. We discretize each pedal cycle into 721 crank positions from *ϕ* = *{* 0°, …, 360°*}*, with 0° at the top and 180° at the bottom dead center. For each crank position *ϕ, L*_c_ defines the pedal position, and the foot geometry defines the ankle position. The anthropometric thigh and shank lengths *L*_thigh_ and *L*_shank_ then determine the position of the knee joint through geometric closure. This geometric representation captures the primary lower-limb degrees of freedom used in contemporary musculoskeletal models of cycling [3]. We select the forward-knee solution and calculate the joint angles throughout the complete pedal cycle. We decompose the hip angle into lower and upper contributions,

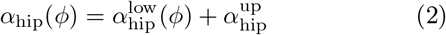

where 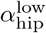 denotes the angle between the knee–hip segment and the horizontal and 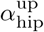 denotes the angle between the horizontal and the hip–shoulder segment. Here, lower and upper refer to the thigh and trunk contributions to the hip geometry. Crank length changes 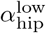 through the lower-limb geometry, whereas torso position determines 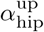. We define the minimum lower hip and knee angles across the pedal cycle as

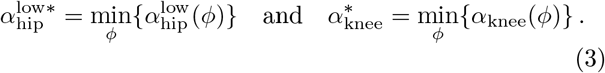

**Figure 1:**
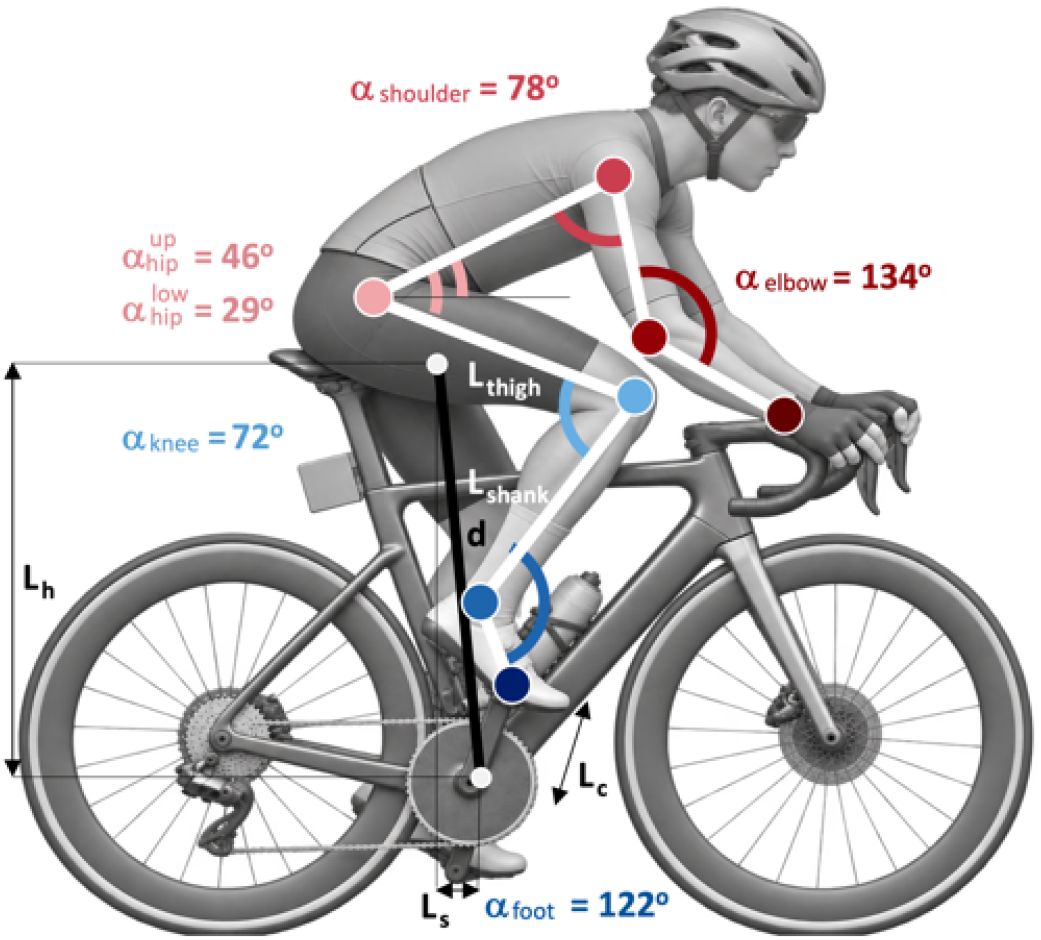
Cycling kinematics and rider geometry. Planar representation of the rider–bicycle system to define the kinematic model. The lower limb consists of thigh and shank segments of lengths *L*_thigh_ and *L*_shank_, connected to the bicycle through the foot. Saddle height *L*_h_ and setback *L*_s_ define the saddle tip position relative to the bottom bracket, with a total distance *d* between both (white circles). The positions of the foot, ankle, knee, hip, shoulder, elbow, and wrist joints characterize the rider configuration and define the joint angles *α*_foot_, *α*_knee_, 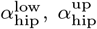, *α*_shoulder_, and *α*_elbow_ (from blue to red). The foot segment is fixed at the end of the crank with crank length *L*_c_ and the hip joint sits at a fixed position above and behind the saddle tip.

We use sex-specific ANSUR II segment proportions to evaluate male and female riders from 150 to 200 cm in stature. To quantify how bicycle geometry affects their hip kinematics, we vary saddle height *L*_h_ from 65 to 75 cm and report their minimum lower hip and knee angles.

### 2.3. Crank length, hip opening, and mechanical power

We quantify how crank length and bicycle geometry affect hip opening and maximal mechanical power. We use the mean male ANSUR II anthropometry and the reference bicycle geometry defined above. First, we fix saddle height *L*_h_ = 70 cm and saddle setback *L*_s_ = 5 cm, vary crank length *L*_c_ = *{* 150, …, 180 *}* mm, calculate the minimum lower hip angle 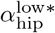, and define the associated hip opening,

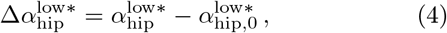

as the increase in minimum lower hip angle relative to the reference configuration. Next, we fix the crank length *L*_c_ = 170 mm, vary saddle height *L*_h_ = *{* 60, …, 80 *}* cm and saddle setback *L*_s_ = *{*−15, …, +5 *}* cm from behind to the front of the bottom bracket and calculate the minimum lower hip angle 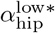 and hip opening 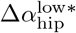. Finally, we quantify the mechanical effect of crank length using the normalized maximal-power relationship [15],

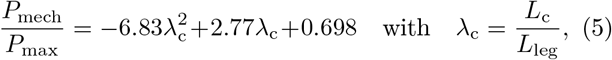

where *L*_leg_ = *L*_thigh_ + *L*_shank_. We scale this relationship by its maximum *P*_max_ and compare against reported maximal-power data [1, 13, 14].

### 2.4. Personalized crank-length analysis

We propagate individual anthropometry through the kinematic model. For all 6,068 individuals with complete stature, thigh, shank, and foot measurements, we scale saddle height *L*_h_ with individual stature *H* as

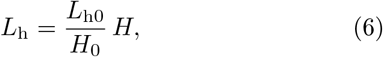

where *L*_h0_ = 70.0 cm is the reference saddle height and *H*_0_ = 175.6 cm is the mean male stature in the database. We use a saddle setback of *L*_s_ = −5 cm for cyclists and a forward saddle position of *L*_s_ = +5 cm for triathletes. For each sex and discipline—male cyclists, female cyclists, male triathletes, and female triathletes—we calculate the reference minimum lower hip angle 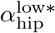 for the mean an-thropometry with a 170 mm crank. For each individual, we then use the measured thigh, shank, and foot lengths, scale saddle height with stature, and solve for the equivalent crank length *L*_c_ that reproduces the group-specific reference minimum lower hip angle. We fit a linear regression between equivalent crank length and stature for each of the four groups and report the scaling coefficient and coefficient of determination *R*^2^. Importantly, this analysis defines equivalent crank lengths for each group’s specific minimum lower hip angles; it does not define or recommend optimal crank lengths.

### 2.5. Elite cyclists and triathletes

To compare the model with crank-length choices in elite sport, we compile a database of publicly available crank-length measurements for professional road cyclists and elite triathletes from 2012 to 2026. For each athlete, we record discipline, sex, stature, crank length, and year. Since individual lower-limb segment lengths are unavailable for the elite athletes, we normalize their observed crank lengths by stature, rather than by leg length. We include road cyclists and triathletes only and exclude other cycling disciplines and temporary experimental crank lengths. For the relationship between crank length and stature, we retain only the most recent configuration per athlete. For temporal analyses, we retain all reported configurations so that repeated observations capture changes within athletes across years. We group the analysis into male cyclists, female cyclists, male triathletes, and female triathletes. Within each group, we perform linear regression to quantify the relationships between crank length and stature, crank length and year, and stature-normalized crank length and year. We report residual standard deviations to quantify the variation around each regression.

### 2.6. Aerodynamic–mechanical power balance

We combine the geometry, aerodynamics, and mechanics into a net-power model that reflects the competition between *aerodynamic* and *mechanical* power changes in response to changes in crank length. Geometrically, shorter crank increases the minimum lower hip angle 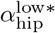. We assume that the rider converts this increase into an equal reduction in upper hip angle while the minimum total hip angle remains constant,

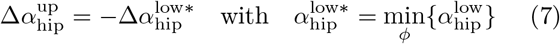

This assumption allows the rider to lower the torso without further closing the hip. *Aerodynamically*, previous studies have shown that, across the range of torso angles considered here, changes in torso angle, or in our model upper hip angle 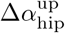, produce approximately linear changes in aerodynamic drag area [17]. We therefore write

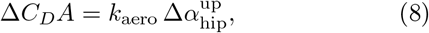

where the aerodynamic sensitivity 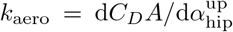 quantifies the change in aerodynamic drag area per degree of upper hip angle. We determine this sensitivity from reported wind-tunnel experiments that measured *C*_*D*_*A* in three competitive cyclists across a range of torso angles at 40 km/h [17], and obtain *k*_aero_ = 0.00277 m^2^*/*deg from linear regression. The resulting change in aerodynamic power is

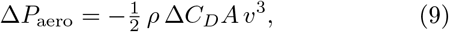

where *v* denotes the cycling speed and *ρ* = 1.225 kg/m^3^ is the air density. *Mechanically*, we adopt the maximal-power relation [15] that introduces changes in mechanical power relative to a reference crank length as

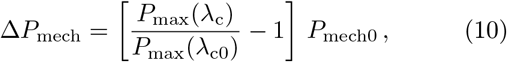

where *P*_mech0_ denotes the reference mechanical power at reference crank length *L*_c0_ = 170 mm, *P*_max_(*λ*_c_) denotes the maximal mechanical power, *λ*_c_ = *L*_c_*/L*_leg_ is the crank-to-leg-length ratio, and *λ*_c0_ = *L*_c0_*/L*_leg_ is the crank-to-leg-length ratio at reference crank length *L*_c0_. A negative value of Δ*P*_mech_ represents a mechanical power loss relative to the reference crank. We define the net power change as the competition between aerodynamic and mechanical effects,

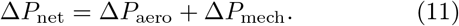

Positive values of Δ*P*_net_ indicate a net power advantage relative to the reference crank, and negative values indicate a net power penalty. A shorter crank provides a net power advantage when the aerodynamic power gain from the lower torso position exceeds the associated mechanical power loss. We evaluate this competition across normalized crank lengths *λ*_c_ = *{* 0.12, …, 0.24 *}* and cycling speeds *υ* = *{* 0, …, 55 *}* km/h to construct sex-specific net-power phase diagrams. We use the corresponding mean ANSUR II anthropometry for each sex and reference mechanical powers of *P*_mech0_ = 300 W for men and *P*_mech0_ = 200 W for women. These reference powers set the *scale* of the mechanical contribution, but do not alter the underlying normalized crank-length dependence of maximal mechanical power. We estimate elite-athlete trajectories for all four groups–men and women, cyclists and triathletes–with representative mechanical powers of *P*_mech0_ = 400 W for men and *P*_mech0_ = 300 W for women. We anchor each trajectory at the observed mean elite normalized crank length and calculate its speed-dependent shift from the model optimum. Importantly, these trajectories combine observed elite crank-length choices with the model prediction for the effect of cycling speed; they do *not* represent crank-length recommendations.

## 3. Results

### 3.1. Population anthropometry

Lower-limb geometry shows systematic sex-specific scaling with stature (Figs. 2 and 3). Across 4,082 men and 1,986 women, mean stature is 175.6 *±* 6.9 cm and 162.8*±*6.4 cm, respectively (Table 1). Mean thigh length is 40.9 *±* 2.8 cm in men and 37.9 *±* 2.4 cm in women, whereas mean shank length is 41.9 *±* 2.5 cm and 40.3 *±* 2.6 cm, respectively (Fig. 2). Segment lengths scale approximately linearly with stature, but the scaling differs between segments and sexes (Fig. 3). Thigh length follows nearly identical proportional scaling in men and women, *λ*_thigh_ = *L*_thigh_*/H* = 0.233 whereas shank length scales as *λ*_shank_ = *L*_shank_*/H* = 0.239 in men and 0.248 in women. Consequently, total thigh-plus-shank length represents approximately 47.2% of stature in men and 48.1% in women. But stature alone does not uniquely determine lower-limb geometry. Individuals near the sex-specific mean stature show substantial variation in both thigh and shank length, and the joint distributions overlap considerably between women and men (Fig. 3, right). Thus, riders of similar stature can have different thigh-to-shank proportions despite the strong population-level scaling of individual segment lengths.

**Table 1:** Population anthropometry. Mean *±* standard deviation for the ANSUR II populations.

| | | men<br>$n=4,082$ | women<br>$n=1,986$ |
| --- | --- | --- | --- |
| stature | $H$ [cm] | $175.6 \pm 6.9$ | $162.8 \pm 6.4$ |
| thigh length | $L_{\text{thigh}}$ [cm] | $40.9 \pm 2.8$ | $37.9 \pm 2.4$ |
| shank length | $L_{\text{shank}}$ [cm] | $41.9 \pm 2.5$ | $40.3 \pm 2.6$ |
| foot length | $L_{\text{foot}}$ [cm] | $27.1 \pm 1.3$ | $24.6 \pm 1.2$ |
| relative thigh | $\lambda_{\text{thigh}}$ [-] | 0.233 | 0.233 |
| relative shank | $\lambda_{\text{shank}}$ [-] | 0.239 | 0.248 |
| relative foot | $\lambda_{\text{foot}}$ [-] | 0.154 | 0.151 |
| relative leg | $\lambda_{\text{leg}}$ [-] | 0.472 | 0.481 |

**Figure 2:**
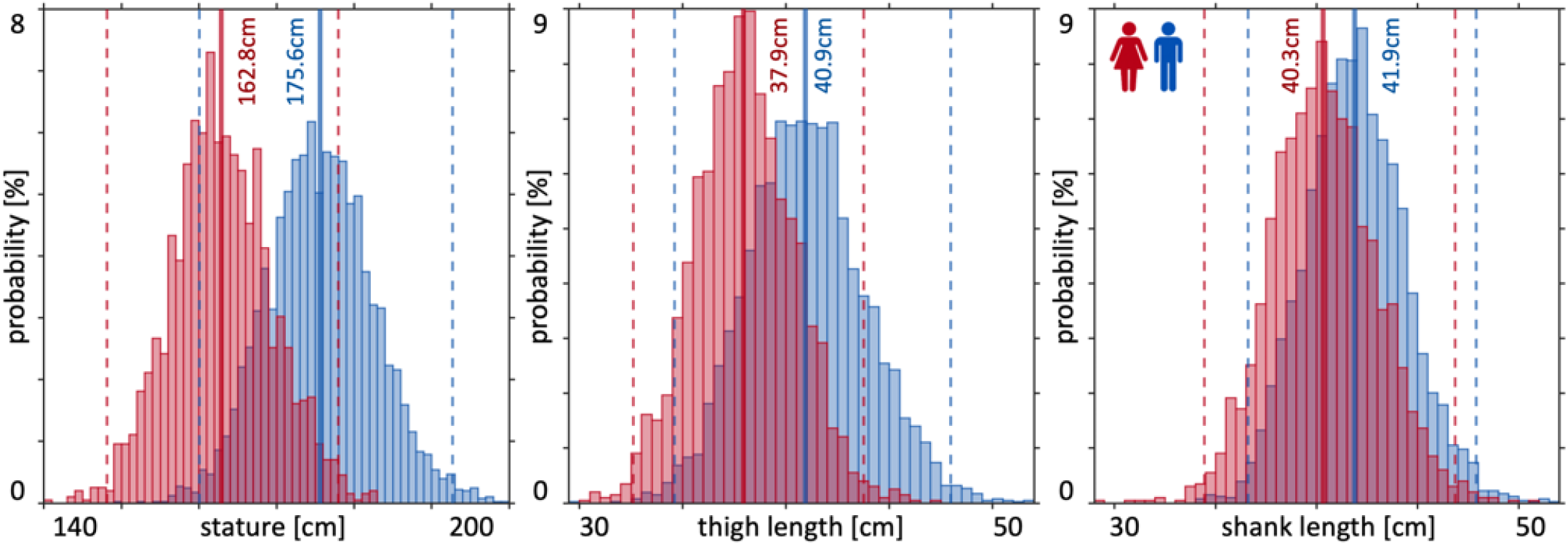
Population distributions of stature and lower-limb segment lengths. Probability distributions of stature (left), thigh length (middle), and shank length (right) for 4,082 men (blue) and 1,986 women (red) from an anthropometric database [8]. Solid lines indicate population means; dashed lines indicate 1st and 99th percentiles.

**Figure 3:**
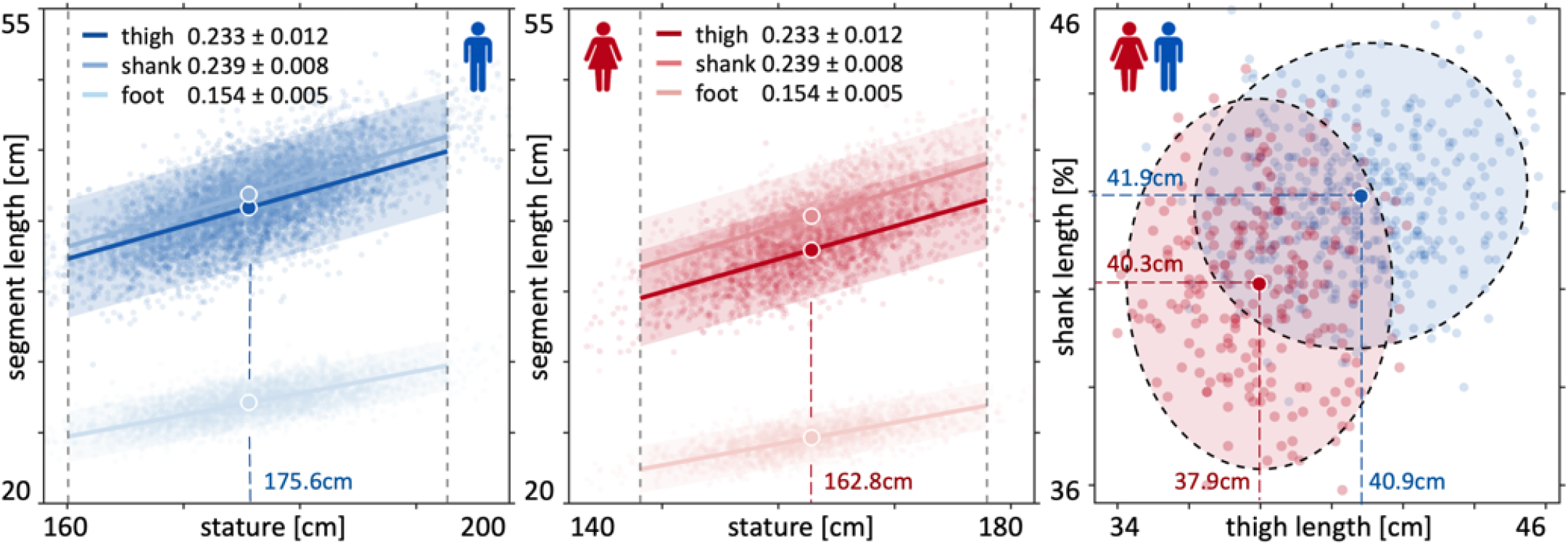
Lower-limb segment lengths scale differently with stature and vary substantially between individuals. Thigh, shank, and foot lengths as functions of stature for 4,082 men (left) and 1,986 women (middle) [8]. Solid lines show population scaling relationships and shaded regions show population variability; circles indicate population means and dashed vertical lines indicate 1st and 99th stature percentiles. Thigh and shank lengths for individuals of similar stature illustrate substantial variation in segment proportions within and between the male and female populations (right). Dashed ellipses indicate joint distributions of thigh and shank length.

### 3.2. Cycling kinematics

Hip and knee angles follow distinct patterns throughout the pedal cycle (Fig. 4). For the reference 170-mm crank, the minimum lower hip angle occurs approximately 45–60° after top dead center across the range of rider statures, whereas the minimum knee angle occurs within approximately 3–5° after top dead center. Interestingly, top dead center does not define the position of maximum hip closure, despite its close correspondence with maximum knee flexion. Rider stature and saddle height systematically shift both joint-angle extrema. At fixed bicycle geometry, taller riders reach smaller minimum lower hip and knee angles, whereas shorter riders maintain more open configurations. Changes in saddle height produce comparable shifts in joint kinematics: higher saddles increase the minimum lower hip and knee angles, whereas lower saddles decrease both angles. Male and female populations follow the same qualitative relationships, but their segment proportions shift the predicted joint angles. For the mean male and female riders at the reference saddle height of 70 cm, the minimum lower hip angles are 29° and 31°, and the minimum knee angles are 72° and 75°. These results establish that crank-induced hip closure emerges from the geometry of the entire lower limb rather than from crank position or stature alone.

**Figure 4:**
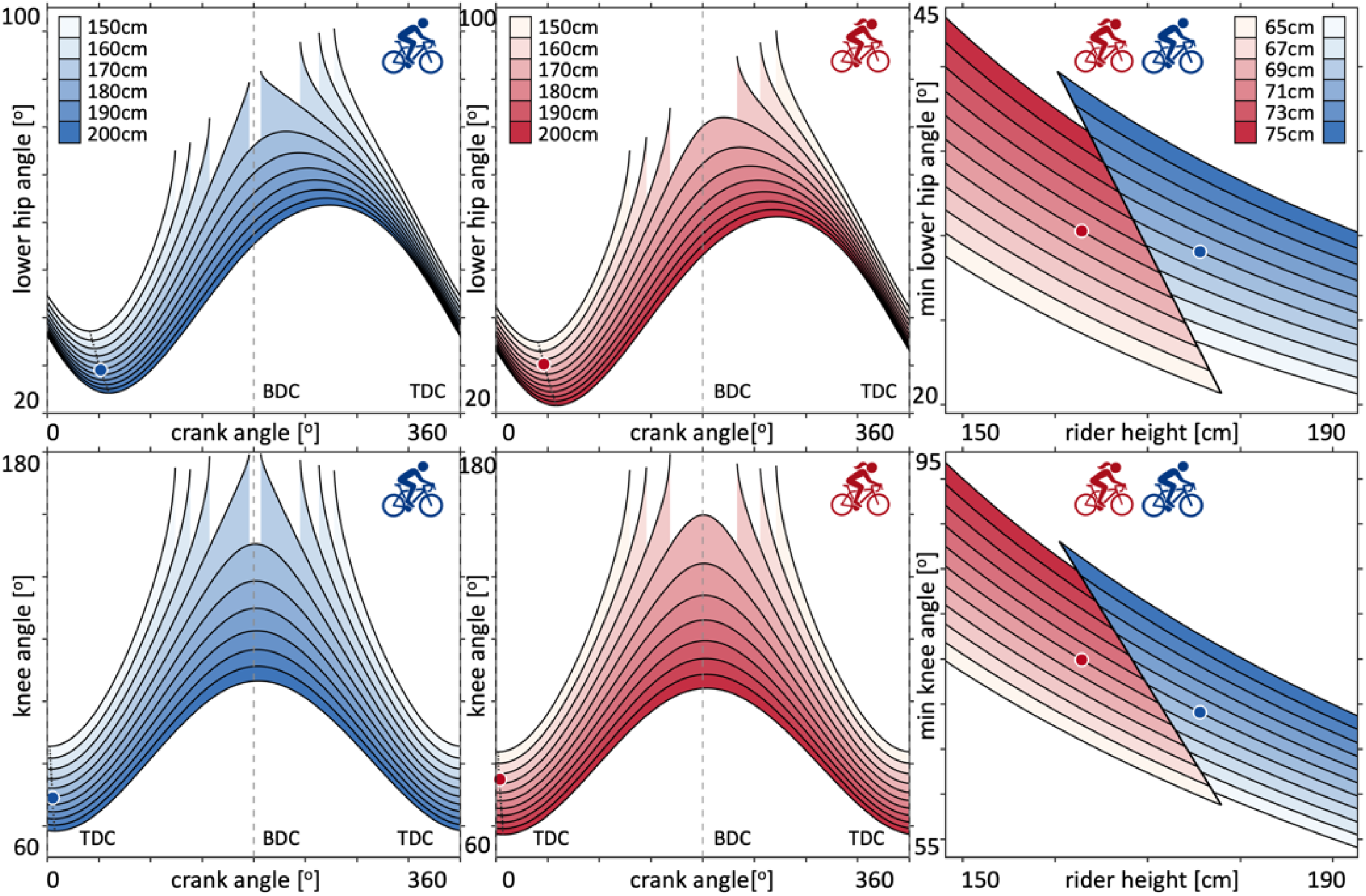
Hip and knee angles follow distinct patterns throughout the pedal cycle. Lower hip angle throughout the pedal cycle for men (top left) and women (top middle) with rider stature from 150 to 200 cm and a 170-mm crank. Minimum lower hip angle occurs after TDC rather than at TDC. Minimum lower hip angle as a function of rider stature and saddle height from 65 to 75 cm (top right). Knee angle throughout the pedal cycle for men (bottom left) and women (bottom middle); minimum and maximum knee angles closely coincide with TDC and BDC. Minimum knee angle as a function of rider stature and saddle height (bottom right). Circles indicate values for mean male and female riders. TDC top dead center, BDC bottom dead center.

### 3.3. Crank length, hip opening, and mechanical power

Crank length directly controls the minimum lower hip angle (Fig. 5, left). Across the practically relevant range from 150 to 180 mm, the nonlinear kinematic solution is nearly perfectly linear,

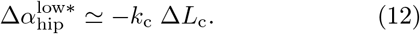

**Figure 5:**
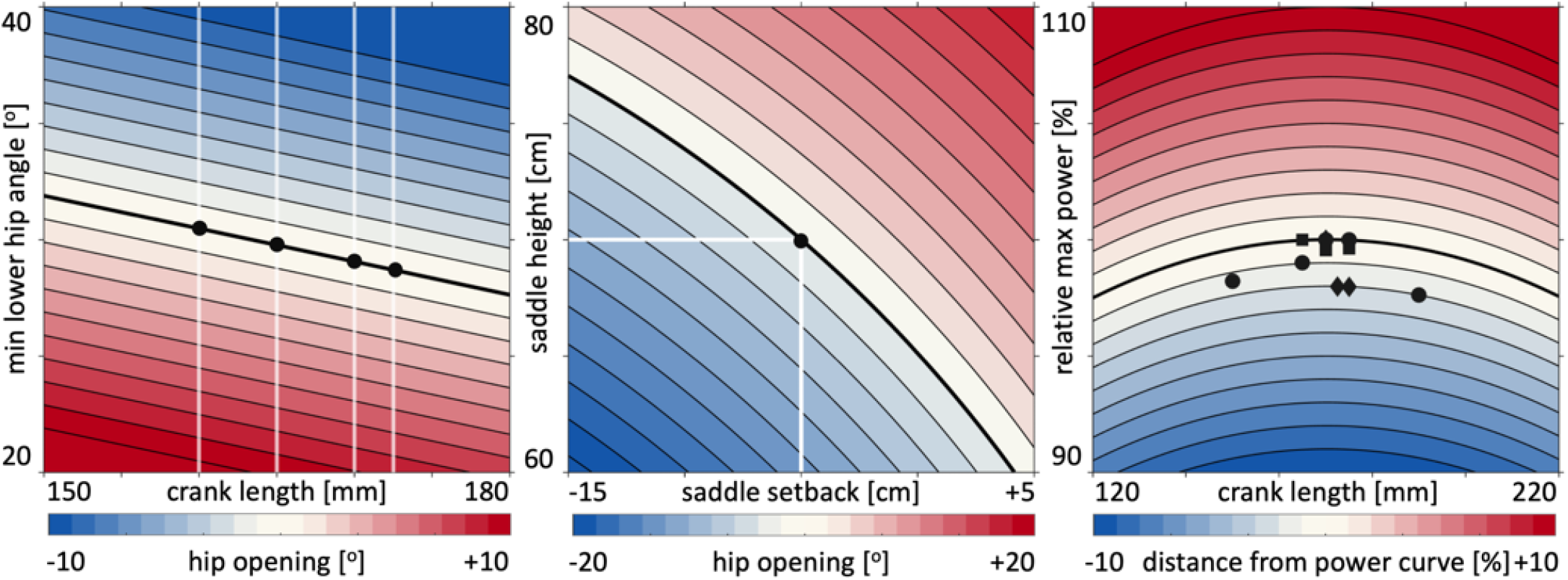
Crank length and bicycle geometry couple hip opening and mechanical power. Minimum lower hip angle across the pedal cycle as a function of crank length (left). Color indicates hip opening relative to the reference position (black), white lines mark common crank lengths. Hip opening at fixed 170 mm crank as a function of saddle height and setback (middle). Color indicates change relative to the reference configuration. Relative maximal power as a function of crank length (right). The black curve shows the mechanical power relation, symbols show experimental data, and color indicates deviation from the maximal-power curve.

For the reference rider, the minimum lower hip angle decreases from 31.9° at 150 mm to 27.6° at 180 mm, and the crank sensitivity 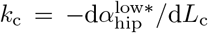 is *k*_c_ = 0.142°*/*mm with *R*^2^ *>* 0.9999. In other words, reducing the crank length *L*_c_ by 10 mm increases the minimum lower hip angle 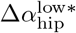 by approximately 1.42°. Bicycle geometry provides an additional mechanism to control hip opening (Fig. 5, middle). Increasing saddle height and moving the saddle forward both increase the minimum lower hip angle, whereas a lower or more rearward saddle closes the hip. Across the investigated range of saddle heights and setbacks, these geometric effects reach several degrees and can exceed the change produced by common differences in crank length. Thus, crank length, saddle height, and saddle setback act together to determine the minimum hip configuration. The crank-length–power relation predicts a broad mechanical optimum near *L*_c_ = 168 mm for the reference rider (Fig. 5, right). Since this optimum scales with leg length, the crank-to-leg-length ratio *λ*_c_ = *L*_c_*/L*_leg_, rather than absolute crank length *L*_c_, provides the appropriate measure for comparisons across riders. Experimental maximal-power measurements follow the same broad response and show only small power differences across commonly used crank lengths. Consequently, shortening the crank within the conventional range produces a measurable increase in hip opening with only a small mechanical-power penalty.

### 3.4. Personalized crank-length analysis

Individual anthropometry produces substantial variation in equivalent crank length at a fixed minimum lower hip angle (Fig. 6). Equivalent crank length increases with stature in all four groups, with scaling coefficients of 0.80 mm/cm for male cyclists, 0.88 mm/cm for female cyclists, 0.42 mm/cm for male triathletes, and 0.49 mm/cm for female triathletes. Thus, the model predicts a stronger stature dependence for the cyclist configuration than for the triathlete configuration. Stature alone, however, explains only a small fraction of the individual variation. Across all four groups, stature explains only 3–13% of the variation in equivalent crank length, with *R*^2^ = 0.13 and 0.09 for male and female cyclists and *R*^2^ = 0.04 and 0.03 for male and female triathletes. Individuals of identical stature can therefore require substantially different crank lengths to preserve the same minimum lower hip angle because their thigh, shank, and foot lengths differ. These results distinguish *population scaling* from *individual prediction*: stature establishes a systematic trend in crank length, but individual lower-limb geometry determines where a rider lies around that trend. The weaker stature dependence in the triathlete configuration further emphasizes that bicycle geometry modifies the relationship between anthropometry and crank length.

**Figure 6:**
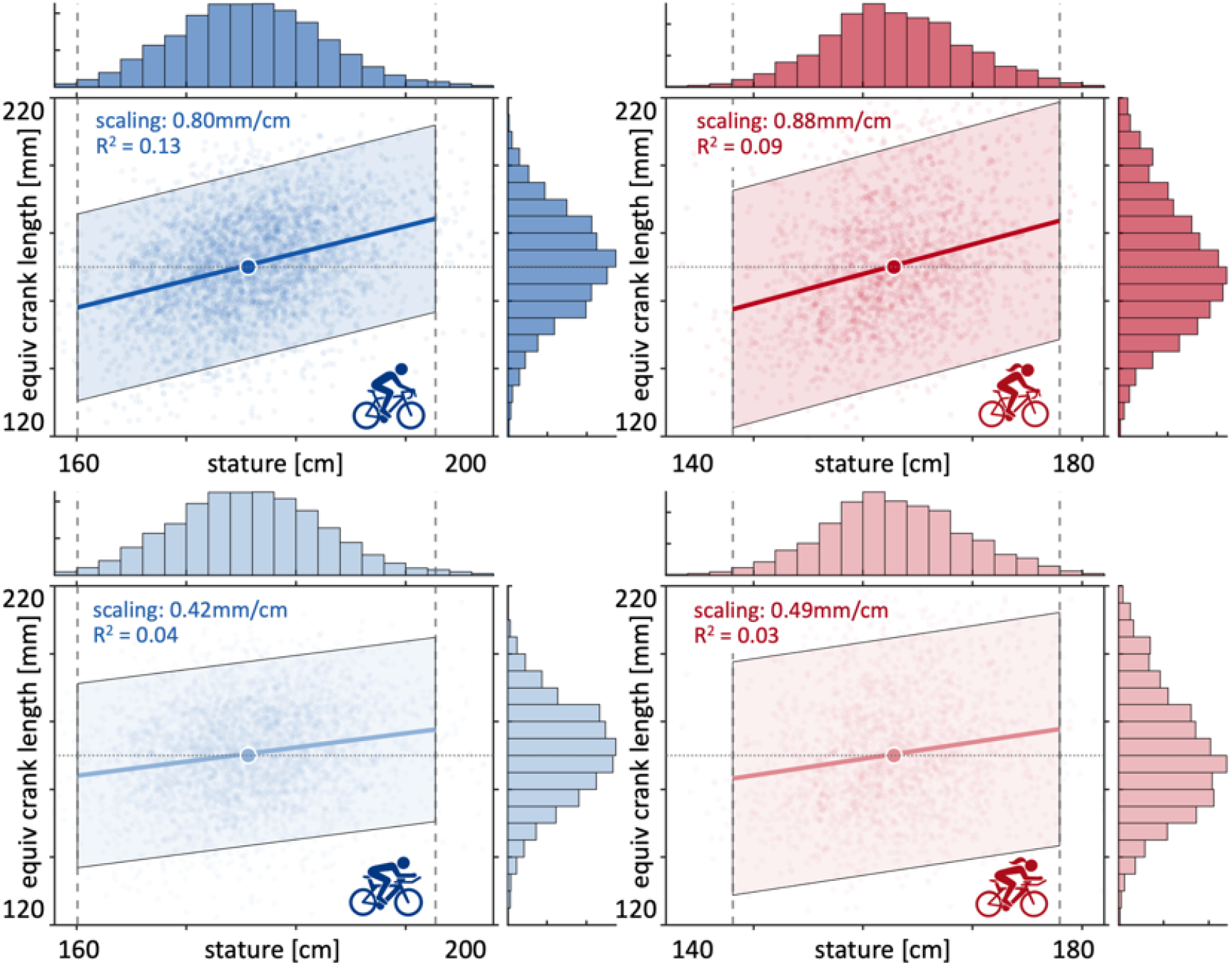
Individual anthropometry produces substantial variation in crank length at fixed lower hip angle. Crank length required to preserve the reference minimum lower hip angle as a function of stature for male cyclists (top left), female cyclists (top right), male triathletes (bottom left), and female triathletes (bottom right). Each point represents an individual from the population with individual thigh, shank, and foot lengths. Solid lines show population scaling relationships; shaded regions show population variability; circles indicate predictions for the mean male and female riders. Marginal histograms show the distributions of stature and predicted crank length. Scaling coe@cients quantify the change in crank length per centimeter of stature; *R*^2^ values quantify the fraction of individual variation explained by stature.

### 3.5. Elite cyclists and triathletes

Crank-length choices show substantial variation across elite athletes (Fig. 7). Across the four elite populations, mean crank length ranges from 164.5 to 170.7 mm, with triathletes using shorter cranks relative to stature than cyclists in both men and women (Table 2). The four current reigning Tour de France winners and Ironman world champions illustrate substantial variation in upper and lower hip angles and overall cycling kinematics at the highest level of the sport (Fig. 7, top). Crank length generally increases with athlete stature, but the large spread within each group confirms that stature alone does not determine crank choice (Fig. 7, left). Consistent with the personalized kinematic analysis (Fig. 6), riders of similar stature can choose substantially different crank lengths. In contrast, crank length shows a clear temporal trend. Elite athletes have progressively adopted shorter cranks over the past decade, with the strongest trends in road cyclists (Fig. 7, middle). Crank length has decreased by approximately 0.61 mm/year in male cyclists (*R*^2^ = 0.43) and 0.38 mm/year in female cyclists (*R*^2^ = 0.31). The broader distributions among triathletes produce weaker temporal relationships, but both male and female triathletes follow the same overall trend toward shorter cranks. Normalization by stature does not eliminate the temporal trend (Fig. 7, right); the progressive reduction in crank length cannot be explained simply by changes in athlete stature over time. Instead, elite practice has shifted toward shorter cranks relative to rider size, consistent with a shift toward crank configurations that permit a lower torso position without further hip closure.

**Table 2:** Elite cyclists and triathletes. Mean *±* standard deviation for the elite athlete populations.

| cyclists | | | men<br>$n=41$ | women<br>$n=35$ |
| --- | --- | --- | --- | --- |
| stature | $H$ | [cm] | $179.4 \pm 7.9$ | $167.5 \pm 6.1$ |
| crank length | $L_c$ | [mm] | $170.7 \pm 4.2$ | $167.4 \pm 3.5$ |
| relative crank | $L_c/H$ | [%] | $9.53 \pm 0.44$ | $10.01 \pm 0.36$ |
| year | | [-] | $2020.4 \pm 4.8$ | $2021.8 \pm 4.8$ |
| triathletes | | | men<br>$n=38$ | women<br>$n=36$ |
| stature | $H$ | [cm] | $182.9 \pm 6.3$ | $169.2 \pm 5.3$ |
| crank length | $L_c$ | [mm] | $168.9 \pm 3.4$ | $164.5 \pm 4.9$ |
| relative crank | $L_c/H$ | [%] | $9.24 \pm 0.31$ | $9.73 \pm 0.24$ |
| year | | [-] | $2021.5 \pm 3.5$ | $2021.7 \pm 3.5$ |

**Figure 7:**
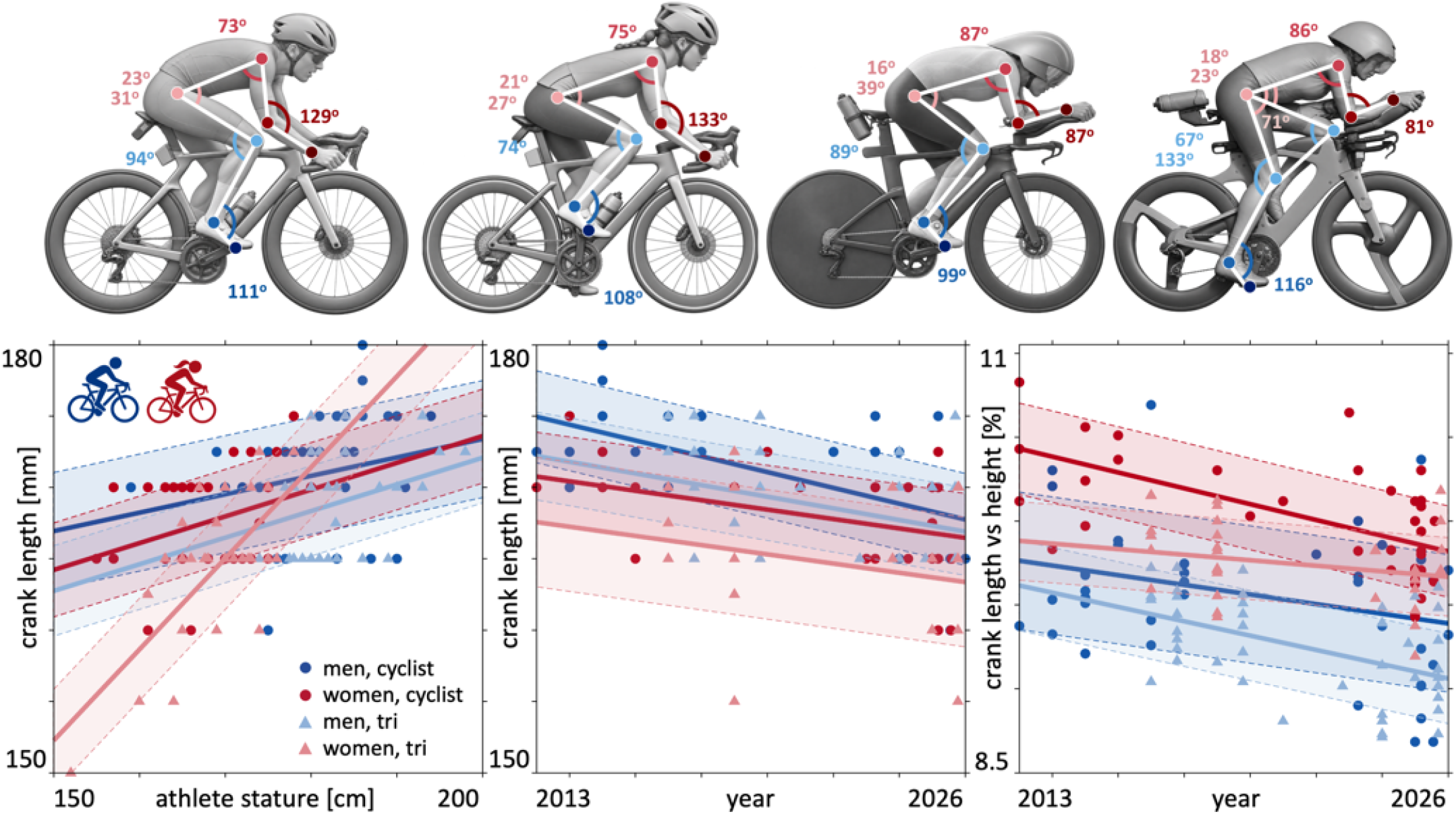
Elite cyclists and triathletes use distinct crank lengths that have shortened over the past decade. Cycling kinematics of the reigning Tour de France winners and Ironman world champions illustrate differences in cycling position across sex and discipline (top). Crank length as a function of athlete stature for male and female cyclists and triathletes (bottom left). Crank length (bottom middle) and crank length normalized by stature (bottom right) as functions in time. Circles denote cyclists and triangles denote triathletes; blue denotes men and red denotes women. Solid lines show group-specific linear regressions and shaded regions show variability.

### 3.6. Aerodynamic–mechanical power model

The competition between mechanical and aerodynamic effects produces a speed dependence of the net-power balance (Fig. 8). A shorter crank increases the minimum lower hip angle and permits a smaller upper hip angle, which reduces aerodynamic drag area. This aerodynamic gain competes with the small mechanical power penalty associated with shorter cranks. At low cycling speeds, crank lengths near the mechanical optimum provide the greatest net-power advantage. As cycling speed increases, the aerodynamic contribution becomes progressively more important: *changes in aerodynamic power scale with the cube of cycling speed*, whereas *changes in mechanical power are independent of speed*. Consequently, the relative importance of the aerodynamic contribution scales as

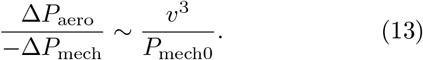

**Figure 8:**
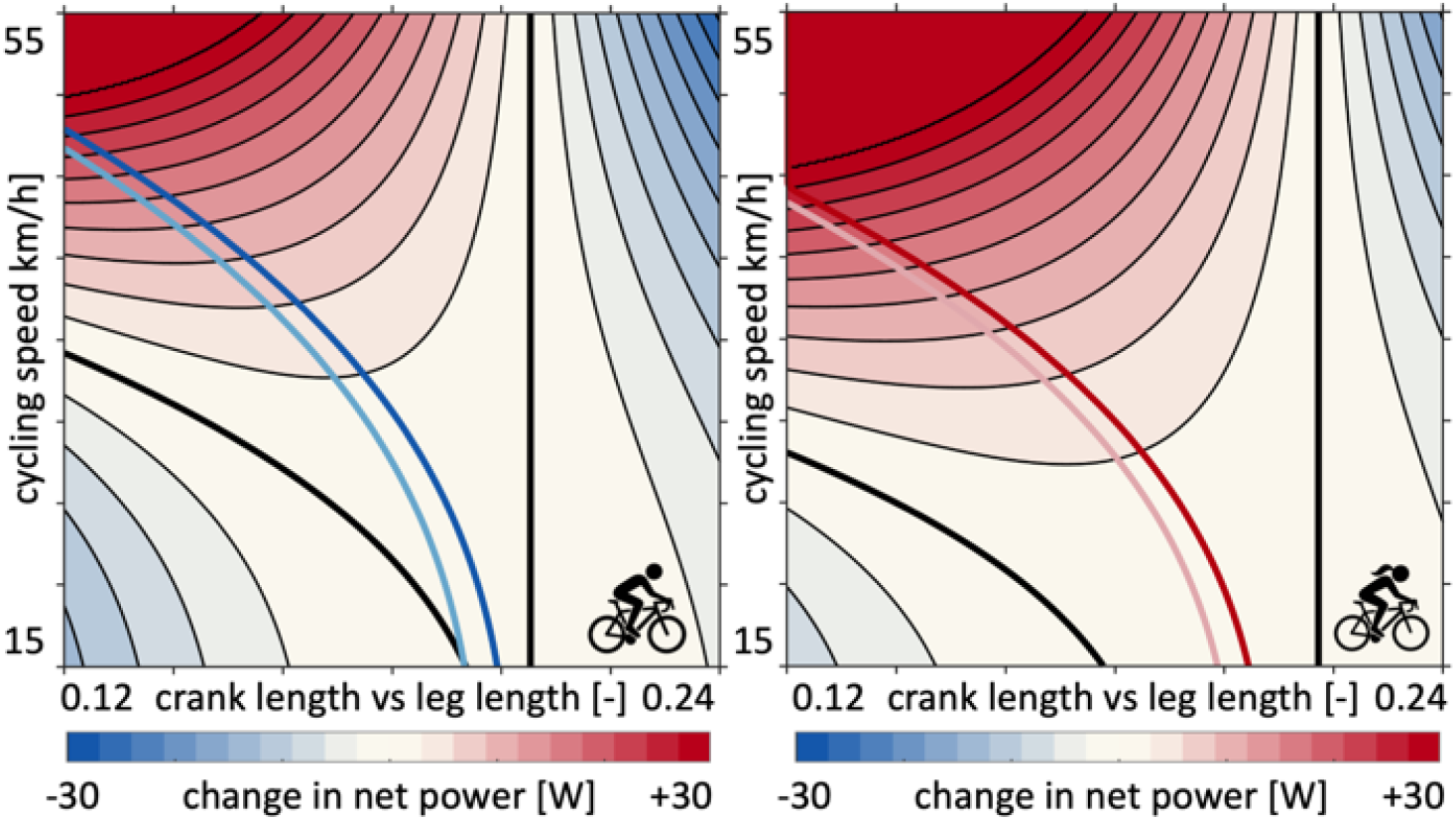
Net-power phase diagrams predict speed-dependent shift toward shorter crank lengths. Change in net power as a function of crank length normalized by leg length and cycling speed for men (left) and women (right). Red regions indicate a net power advantage and blue regions a net power disadvantage relative to the reference crank length; thick black lines indicate zero net power change. Dark and light trajectories show the predicted crank-length scaling for cyclists and triathletes, anchored to the corresponding elite-athlete populations. Vertical black lines indicate the normalized reference crank length.

Higher reference mechanical power *P*_mech0_ shifts the predicted trajectories toward longer crank lengths, whereas higher cycling speed *v* shifts them toward shorter crank lengths. The black zero-net-power contours show this speed-dependent shift toward shorter normalized crank lengths in both men and women. The elite-athlete trajectories follow the same pattern. Differences between the cyclist and triathlete trajectories arise from their population-specific crank lengths and bicycle configurations. Together, these results identify *cycling speed* as the key variable that governs the balance between mechanical and aerodynamic crank-length effects.

## 4. Discussion

Our results reveal a simple but previously underappreciated role of crank length in cycling: A shorter crank opens the hip angle, permits a lower torso position, reduces aerodynamic drag, and enables faster cycling. Crank length emerges not simply as an anthropometric or mechanical parameter, but as a personalized design variable that couples rider geometry to aerodynamics.

### Limitations

Our attempt is *not* to provide a design tool for crank-length selection and has a few limitations. First, we keep saddle height and setback fixed when crank length changes, although riders may adjust both in practice. Second, we prescribe foot angle throughout the pedal cycle; a more refined model could determine foot motion from individual cycling kinematics. Third, our net-power model isolates crank-dependent mechanical and aerodynamic effects and does not include other contributions to cycling power, such as rolling and drivetrain resistance. Fourth, we assume that the hip-angle margin created by shorter cranks can translate directly into a lower torso position and estimate the resulting change in aerodynamic drag area from published wind-tunnel data. These assumptions deliberately reduce the complexity of the rider–bicycle system to expose the global relationships between crank length, geometry, and aerodynamics and identify opportunities for future model refinement.

### Stature poorly predicts personalized crank length

Population anthropometry reveals systematic sex- and stature-dependent differences in lower-limb geometry, but *stature alone* poorly predicts individualized crank length. Leg length averages 47.2% of stature in men and 48.1% in women, yet individuals of similar stature show substantial variation in thigh and shank proportions. This variability propagates directly into cycling kinematics: across our four sex–discipline groups, stature explains only 3–13% of the variation in the crank length that preserves the reference minimum lower hip angle. Previous experiments similarly found that crank-to-leg-length and crank-to-tibia-length ratios explain only 20.5% and 21.1% of the variability in maximal cycling power, despite a population-level power optimum near 20% of leg length [15]. Our results extend this observation from power to geometry. Sex-specific scaling captures systematic population differences, but individual thigh, shank, and foot lengths determine where a rider lies within those distributions [10]. A universal crank-to-stature ratio therefore misses the *individual geometry* that governs hip closure.

### Crank length controls geometry more than power

Across the conventional range, crank length has a pronounced effect on rider geometry but a remarkably small effect on mechanical power. Maximal power varies by only about 4% across the extreme range from 120 to 220 mm [15], and joint-specific power changes little across 150–190 mm at each rider’s optimal pedal rate [1]. Under sub-maximal conditions, *±*5-mm crank changes produce no significant change in gross efficiency but alter maximum hip and knee flexion and range of motion by 1.8–3.4° [5]. Recent experiments confirm this separation: crank lengths from 165 to 175 mm produce no significant differences in cycling efficiency or sprint power in trained cyclists [13], and reductions of 5–10 mm reduce hip and knee flexion without a detectable change in mean power [16]. Our model explains and quantifies these observations: each 1-mm crank reduction increases the minimum lower hip angle by 0.142°, so a 10-mm reduction provides approximately 1.42° of additional hip opening. This simple scaling reframes the mechanically neutral regime as *geometric freedom*: riders can alter crank length at little mechanical cost but with a predictable change in hip closure.

### Speed amplifies the aerodynamic value of geometry

A shorter crank can preserve hip opening at a lower torso position, and wind-tunnel experiments show that torso angle strongly affects aerodynamic drag area [17]. Across the range relevant here, these measurements give *k*_aero_ = 0.00277 m^2^*/*deg. Together, our two sensitivities predict that a 10-mm crank reduction can support a 1.42° torso-angle change and a 0.00393 m^2^ reduction in *C*_*D*_*A*. The corresponding aerodynamic advantage rises from 3.30 W at 40 km/h to 4.71 W at 45 km/h because aerodynamic power scales with *v*^3^. Previous time-trial models independently predict progressively lower optimal torso angles as speed rises and place the crossover at which aerodynamic losses exceed physiological power losses near 46 km/h [6]. Field [9], wind-tunnel [7], and computational [4] studies independently show that rider position strongly affects aerodynamic drag. Our analysis provides the missing link between these bodies of literature: *crank length couples lower-limb geometry to the speed-dependent aerodynamic trade-ofl*.

### Triathlon magnifies small aerodynamic gains

Previous work shows that the optimal time-trial position reflects the competition between aerodynamic drag and physiological power [6], whereas systematic evidence provides no universal recommendation for crank length [11]. Contemporary elite practice follows the direction that our model predicts. Both 2025 Ironman world champions, Casper Stornes and Solveig Løvseth, raced on 165-mm cranks, consistent with the recent shift toward shorter cranks in elite triathlon. The consequences can become substantial over the 180-km non-drafting bike leg, where athletes maintain an aerodynamic position for several hours. A striking historical comparison comes from the women’s Ironman World Championship in Kona. Daniela Ryf set the current 4:26:07 bike course record in 2018 on 172.5-mm cranks, whereas 2025 world champion Solveig Løvseth raced on 165-mm cranks. Our scaling law predicts that the same 7.5-mm reduction would open the lower hip angle by approximately 1.07°. If a rider converts this geometric freedom entirely into a lower torso position, our aerodynamic scaling predicts a reduction in drag area of Δ*C*_*D*_*A* = −0.00295 m^2^, equivalent to approximately 2.6 W at the current women’s course record pace. A first-order estimate that considers aerodynamic drag alone predicts that this reduction would lower her 4:26:07 bike split by approximately 1:09 min. This thought experiment does not imply that shorter cranks would necessarily have made Daniela Ryf faster; it isolates the potential aerodynamic value of crank-mediated geometric freedom. Our model therefore does not prescribe a universal crank length. Instead, it reveals why *shorter cranks can translate geometric freedom into meaningful performance gains* when speed is high, drafting is absent, and duration is long.

## Conclusion

Crank length has traditionally been viewed as an anthropometric or mechanical design variable, yet mechanical power and cycling economy remain remarkably insensitive to crank length across a broad range. Here we show that its more important role may be geometric: Population anthropometry and cycling kinematics reveal that stature alone poorly predicts personalized crank length, while shorter cranks systematically increase hip-angle margin with little mechanical penalty. This geometric freedom permits a lower aerodynamic position, and its aerodynamic value grows rapidly with cycling speed because aerodynamic power scales with the velocity cubed. Elite cycling and triathlon data support this shift toward shorter cranks, particularly in disciplines where aerodynamic drag is a major determinant of cycling performance. Together, these results establish crank length as a personalized design variable that couples lower-limb geometry to aerodynamics. Rather than asking which crank length maximizes power, future bicycle optimization should ask which crank length best integrates rider geometry, aerodynamic position, and performance objectives. This framework provides a path toward personalized crank selection, integrated rider–bicycle design, and performance optimization for the increasing aerodynamic demands of high-speed cycling.

## Acknowledgements

The authors acknowledge inspiration from the Global Cycling and Global Triathlon Networks GCN and GTN, and support from the Wu Tsai Human Performance Alliance, the NSF CMMI grant 2320933, and the ERC Advanced Grant 101141626.

## CRediT authorship contribution statement

Scott Delp: Conceptualization, Methodology, Review and Editing. Ellen Kuhl: Conceptualization, Methodology, Software, Formal analysis, Data Curation, Validation, Writing Original Draft, Review and Editing.

## Data availability

Data and source code are available at https://github.com/LivingMatterLab.

## Statement of AI-assisted tools usage

This document was prepared with the assistance of OpenAI’s ChatGPT. The authors reviewed, edited, and take full responsibility for the content and conclusions of this work.

## References

[1] Barratt, P.R., Kor”, T., Elmer, S.J., Martin, J.C., 2011. Effect of crank length on joint-specific power during maximal cycling. Med. Sci. Sports Exerc. 43, 1689–1697.

[2] Barratt, P.R., Martin, J.C., Elmer, S.J., Korff, T., 2016. Effects of pedal speed and crank length on pedaling mechanics during submaximal cycling. Med. Sci. Sports Exerc. 48, 705–713.

[3] Clancy, C.E., Gatti, A.A., Ong, C.F., Maly, M.R., Delp, S.L., 2023. Muscle-driven simulations and experimental data of cycling. Sci. Rep. 13, 21534.

[4] Defraeye, T., Blocken, B., Koninckx, E., Hespel, P., Carmeliet, J., 2010. Aerodynamic study of different cyclist positions: CFD analysis and full-scale wind-tunnel tests. J. Biomech. 43, 1262–1268.

[5] Ferrer-Roca, V., Rivero-Palomo, V., Ogueta-Alday, A., Rodríguez-Marroyo, J.A., García-López, J., 2017. Acute effects of small changes in crank length on gross e!ciency and pedalling technique during submaximal cycling. J. Sports Sci. 35, 1328–1335.

[6] Fintelman, D.M., Sterling, M., Hemida, H., Li, F.-X., 2014. Optimal cycling time trial position models: Aerodynamics versus power output and metabolic energy. J. Biomech. 47, 1894–1898.

[7] García-López, J., Rodríguez-Marroyo, J.A., Juneau, C.-E., Peleteiro, J., Córdova Martínez, A., Villa, J.G., 2008. Reference values and improvement of aerodynamic drag in professional cyclists. J. Sports Sci. 26, 277–286.

[8] Gordon, C.C., Blackwell, C.L., Bradtmiller, B., Parham, J.L., Barrientos, P., Paquette, S.P., Corner, B.D., Carson, J.M., Venezia, J.C., Rockwell, B.M., Mucher, M., Kristensen, S., 2014. 2012 Anthropometric Survey of U.S. Army Personnel: Methods and Summary Statistics. Technical Report NATICK/TR-15/007, U.S. Army Natick Soldier Research, Development and Engineering Center, Natick, MA.

[9] Grappe, F., Candau, R., Belli, A., Rouillon, J.D., 1997. Aerodynamic drag in field cycling with special reference to the Obree’s position. Ergonomics 40, 1299–1311.

[10] Hull, M.L., Gonzalez, H., 1988. Bivariate optimization of pedalling rate and crank arm length in cycling. J. Biomech. 21, 839–849.

[11] Husband, S.P., Wainwright, B., Wilson, F., Crump, D., Mockler, D., Carragher, P., Nugent, F., Simms, C.K., 2024. Cycling position optimisation—a systematic review of the impact of positional changes on biomechanical and physiological factors in cycling. J. Sports Sci. 42, 1477–1490.

[12] Inbar, O., Dotan, R., Trousil, T., Dvir, Z., 1983. The effect of bicycle crank-length variation upon power performance. Ergonomics 26, 1139–1146.

[13] Li, J., Wang, Q., Zhang, Y., Zhang, L., Dong, G., Ma, G., Peng, S., Wang, B., Wang, J., Zhou, J., Bao, D., 2025. Effects of crank length on cycling e!ciency, sprint performance, and perceived fatigue in high-level amateur road cyclists. J. Exerc. Sci. Fit. 23, 175–180.

[14] MacDermid, P.W., Edwards, A.M., 2010. Influence of crank length on cycle ergometry performance of well-trained female cross-country mountain bike athletes. Eur. J. Appl. Physiol. 108, 177–182.

[15] Martin, J.C., Spirduso, W.W., 2001. Determinants of maximal cycling power: crank length, pedaling rate and pedal speed. Eur. J. Appl. Physiol. 84, 413–418.

[16] Reynolds, S., Chidley, J., Briley, S., Outram, T., 2026. The impact of minor crank length adjustments on lower body cycling kinematics. Sports Biomech. 25, 454–467.

[17] Underwood, L., Schumacher, J., Burette-Pommay, J., Jermy, M., 2011. Aerodynamic drag and biomechanical power of a track cyclist as a function of shoulder and torso angles. Sports Eng. 14, 147–154.

